# Co-invasion of the ladybird *Harmonia axyridis* and its parasites *Hesperomyces virescens* fungus and *Parasitylenchus bifurcatus* nematode to the Caucasus

**DOI:** 10.1101/390898

**Authors:** Marina J. Orlova-Bienkowskaja, Sergei E. Spiridonov, Natalia N. Butorina, Andrzej O. Bieńkowski

## Abstract

Study of parasites in recently established populations of invasive species can shed lite on sources of invasion and possible indirect interactions of the alien species with native ones. We studied parasites of the global invader *Harmonia axyridis* (Coleoptera: Coccinellidae) in the Caucasus. In 2012 the first established population of *H. axyridis* was recorded in the Caucasus in Sochi (south of European Russia, Black sea coast). By 2018 the ladybird has spread to the vast territory: Armenia, Georgia and south Russia: Adygea, Krasnodar territory, Stavropol territory, Dagestan, Kabardino-Balkaria and North Ossetia. Examination of 213 adults collected in Sochi in 2018 have shown that 53% of them are infested with *Hesperomyces virescens* fungi (Ascomycota: Laboulbeniales) and 8% with *Parasitylenchus bifurcatus* nematodes (Nematoda: Tylenchida, Allantonematidae). Examined *H. axyridis* specimens were free of parasitic mite *Coccipolipus hippodamiae*. An analysis of the phylogenetic relationships of *Parasitylenchus bifurcatus* based on 18S rDNA confirmed the morphological identification of this species. *Hesperomyces virescens* and *Parasitylenchus bifurcatus* are firstly recorded from the Caucasus and Russia, though widespread in Europe. It probably indicates that they appeared as a result of co-invasion with their host. *Harmonia axyridis* was released in the region for pest control, but laboratory cultures are always free of *H. virescens* and *P. bifurcatus*. Therefore, detection of *H. virescens* and *P. bifurcatus* indicates that population of *H. axyridis* in the Caucasus cannot derive exclusively from released specimens. We did not find *H. virescens* on 400 specimens of 31 other ladybird species collected in the same localities with *H. axyridis* in the Caucasus. No reliable correlation between infestation by *H. virescens* and *P. bifurcatus* has been found. Besides these two parasites an unidentified species of the order Mermithida is recorded. It is the first documented case of *H. axyridis* infestation by a parasitic nematode of this order in nature.

## Introduction

Despite a large body of work on invasion ecology, interactions of invasive species with their natural enemies, in particular parasites, are poorly studied [1]. Study of parasites of alien species in young, recently established populations is of great importance for understanding of rotes of invasion [2]. Besides that, some parasites of alien species can affect native species [3]. Therefore, the study of parasites might reveal possible indirect interactions of an alien species with native ones and some reasons of its invasive success. The aim of our investigation was to determine what parasites affect the harlequin ladybird *Harmonia axyridis* (Pallas) (Coleoptera: Coccinellidae) in the Caucasus recently invaded by this species.

*Harmonia axyridis* native to East Asia has been introduced widely for biological control of agricultural pests and established almost all over the world (see detailed description of the native range [4] and overview of global invasion [5]). It established in Western Europe the late 1990s and then expanded its range rapidly. Outbreak of *H. axyridis* in some regions caused a number of negative ecological consequences including decline of native ladybird species [6]. Approximately by 2010 the expansion of European range to the east reached Russia and adjacent regions [7]. The reasons of the great invasive success of this species and decline of native ladybirds attract attention of hundreds of scientists (see review by Roy et al. [5]). Recent studies have shown that some symbionts of this global invader (parasites and microorganisms) have contributed to its success and sometimes even become a “biological weapon” against its competitors - native ladybirds [1, 3, 8].

*Harmonia axyridis* was widely used for biological control of Aphidae and other pests in the Caucasus since 1927. In particular, in the 1980s more than 107,000 of specimens brought from the Far East were released in Georgia [9]. But in spite of these massive releases, *H. axyridis* did not establish before the 21st century. The first established population in the region was recorded in Sochi in 2012 [10, 11]. Then *H. axyridis* quickly spread and became common all over the Black Sea coast of the Caucasus and in adjacent regions. By 2018 it has been recorded in Armenia, Georgia including Abkhazia, and south Russia: Adygea, Krasnodar territory, Stavropol territory, Rostov region, Dagestan, Kabardino-Balkaria and North Ossetia (Fig 1) [12–14]. The releases of *H. axyridis* continued at least to 2010 [15]. So it was unclear if the population in the Caucasus originated from released specimens or appeared a result of expansion of European invasive range of the species [11, 14, 16].

**Fig 1.**
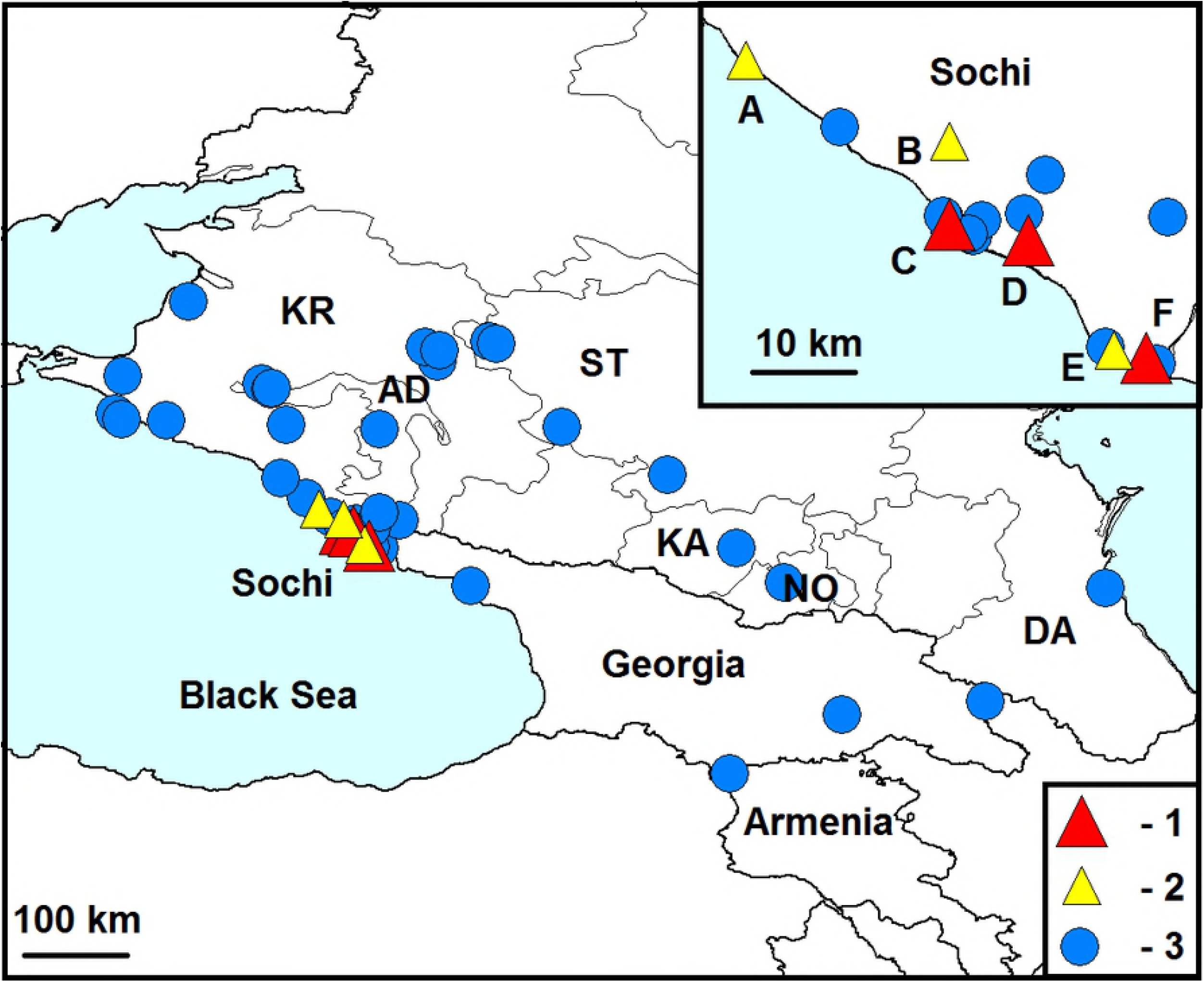
Known localities of *Harmonia axyridis* and its parasites in the Caucasus.

1 – localities of infestation of *Harmonia axyridis* with *Hesperomyces virescens* and *Parasitylenchus bifurcatus*, 2 – localities of infestation of *Harmonia axyridis* with *Hesperomyces virescens*, 3 – other localities of *Harmonia axyridis*. Regions of Russia: AD – Adygea, DA – Dagestan, KA – Kabardino Balkaria, KR – Krasnodar region, NO – North Ossetia, ST – Stavropol region. Localities of detection of the parasites: A – Golovinka, B – Razbityj Kotel, C – Central District, D – Agur, E – Adler, F – Veseloe. Description of localities and sources of information are indicated in supporting information (S1 Table). The map is compiled with the help of DIVA GIS program using the free basic map: http://diva-gis.org/Data.

Establishment of *H. axyridis* in the Caucasus causes three main questions:

1. What are the sources of invasion of *H. axyridis* to the Caucasus?
2. Why is *H. axyridis* currently established in the region, though previously failed to establish in spite of massive releases?
3. How will *H. axyridis* affect native ladybird species?

Though our study of parasites cannot directly answer these difficult questions, it could probably shed some light on them.

We restrict usage of the term ‘‘parasites’’ to those organisms living at the expense of a single host, which are multicellular (in contrast to pathogenic microorganisms) and do not directly cause death of the host (in contrast to parasitoids) [1]. Three species of parasites have been recorded to infest *Harmonia axyridis* in the world: *Hesperomyces virescens* Thaxt. fungi, *Coccipolipus hippodamiae* (McDaniel and Morrill) mites, and *Parasitylenchus bifurcatus* Poinar and Steenberg nematodes [1].

*Hesperomyces virescens* (Ascomycota: Laboulbeniales) is an obligate ectoparasite that has been reported to infect adults of over 30 ladybird species (Coleoptera: Coccinellidae) in all continents except Australia and Antarctica [1]. The association *H. axyridis–H. virescens* has been reported from Austria, Belgium, Czech Republic, Croatia, Germany, France, Hungary, the Netherlands, Poland, Slovakia, the UK, USA, Canada, Argentina, Ecuador, South Africa, and China [1]. *Hesperomyces virescens* has not been detected in Russia or in the Caucasus before [1]. Native range of this fungus is unknown. It was assumed that it could originate from North America [17]. Its entire life cycle takes place on integument of adult ladybird. The sticky spores have shorts life span, are exclusively spread by activities of the host, and transmission takes place during direct contact of specimens (mating or overwintering in aggregations) [18].

*Coccipolipus hippodamiae* (Acarina: Podapolipidae) is an ectoparasitic mite. All life stages live on the underside of the elytra of an adult ladybird and feed on the haemolymph. It is reported to affect five Coccinellidae species in North America, Africa and Europe. Transmission also takes place during mating or overwintering in aggregations.

*Parasitylenchus bifurcatus* (Nematoda: Allantonematidae) is an obligate endoparasite. Its only known host is *Harmonia axyridis*. Fertilized infective female enters an adult ladybird through spiracles or thin parts of the integument. It develops into a female of the first parasitic generation, producing eggs, from which the second parasitic generation females develop. The latter give birth to juvenile males and females, maturing and mating inside the body of the host. Then new infective females appear and leave the host. Mechanism of transmission is unknown [19]. *Parasitylenchus bifurcatus* originally described from Denmark [19], has been also reported in the Netherlands [20], Czech Republic, Poland [1], the USA and Slovenia [21]. There have been no documented records of infestation of *H. axyridis* by nematodes belonging to the order Mermithida in nature.

No parasites of *H. axyridis* in the Caucasus and European Russia have been recorded [5]. We have found *Hesperomyces virescens*, *Parasitylenchus bifurcatus* and an unidentified species belonging to the order Mermithida on *H. axyridis* in the region.

## Materials and methods

### Collection of Harmonia axyridis

Adult *Harmonia axyridis* (213 specimens) were collected in the city of Sochi (south of European Russia, Krasnodar region, Black sea coast of the Caucasus) in April 17–May 15 2018. Localities of collection: Veseloe (43.41, 39.98), Central District (43.58, 39.73), Adler (43.42, 39.94), valley of Agur river (43.56, 39.83), Golovinka (43.79, 39.47), Razbityj Kotel (43.69, 39.73). The beetles were collected by shaking of different trees and shrubs branches and sweep-netting of grasses and collected by hand. Probably all these specimens have overwintered, since the larvae were firstly detected on May 1 and pupae on May 11. No specimens of young generation with soft elytra were detected. All beetles were placed in plastic containers and kept alive at a temperature of about +4°C.

### Screening for parasites

In mid-June each specimen was examined under a stereomicroscope to detect if it was infested with parasites. First, the ladybird was examined externally from above and from below to determine the presence of ectoparasites. Then its elytra were removed to detect if tergites were infested with mites or other parasites. Then abdomen of each specimen was dissected to find endoparasites. This method allows to detect different parasites in one specimen. Dissection was performed in 0.9% NaCl solution. Collected nematodes were heat-killed (at 65°C) and fixed with TAF solution. Permanent slides of nematodes in anhydrous glycerin were prepared following the Seinhorst method [22]. Totally 60 permanent slides were made. Permanent nematode slides and TAF-fixed nematodes are kept in the Helminthological Museum of the Russian Academy of Sciences (Moscow). Morphometric analysis encompassing measurements of common nematode body features was carried out on five fixed nematode specimens of each life stage, employing Zeiss Jenaval microscope. Photographs were taken employing Leica 5500B microscope.

### Identification

Primarily the identification of the parasites was based on morphological features. Identification of *Parasitylenchus bifurcatus* was confirmed through nucleotide sequences analysis. The nematodes recovered from the ladybird haemocoel were frozen individually in sterile 0.7 ml Eppendorf tubes for DNA extraction, which was performed according to Holterman *et al*. [23]. The worm-lysis solution (950 μl of a mixture of 2 ml of 1M NaCl, 2 ml of 1M Tris-HCl, pH 8 plus 5.5 ml of deionised water plus 10 μl of mercaptoethanol and 40 μl of proteinase K, 20 mg ml^−1^) was prepared directly before DNA extraction. Aliquots of 25 μl of sterile water and 25 μl of worm-lysis solution were added to each tube with a nematode and incubated at 65°C for 90 min. The tubes containing the homogenate were then incubated at 99°C for 5 min to deactivate proteinase K. About 1.0 μl of homogenate was used as PCR template.

PCR reactions were performed using Encyclo Plus PCR kit (Evrogen®, Moscow, Russia) according to the manufacturer’s protocol. Primer pairs Nem18S_F (5’-CGC GAA TRG CTC ATT A CA ACA GC-3’) and 26R (5’-CAT TCT TGG CAA ATG CTT TCG-3’) were used to obtain partial (about 900 bp long) sequence of 5’ half of the mitochondrial 18S rDNA [24]. PCR cycling parameters included primary denaturation at 94°C for 5 min followed by 34 cycles 94°C for 45 s, 54°C for 60 s and 72°C for 1 min, followed by post-amplification extension at 72°C for 3 min.

A pair of primers D2A (5’-ACA AGT ACC GTG AGG GAA AGT TG -3’) and D3B (5’-TCG GAA GGA ACC AGC TAC TA-3’) was used to amplify approx. 800 bp long sequence of D2D3 expansion segment of 28S rDNA [25]. PCR cycling parameters included denaturation at 95°C for 3 min, followed by 35 cycles of 94°C for 30 s, 54°C for 35 s, and 72°C for 70 s and followed by post-amplification extension at 72°C for 5 min.

PCR products were visualised in 1% agarose gel. Then bands containing all obtained PCR product were excised from 0.8% agarose gel for DNA extraction with Wizard SV Gel and PCR Clean-Up System (Promega, Madison, USA). Samples were directly sequenced using the same primers as used for primary PCR. The sequences were combined and aligned using the Clustal_X program after the addition of sequences from the GenBank [26]. Similar sequences were searched for in NCBI GenBank with BLAST algorithm [27]. Subsequently, the sequences were edited using the Genedoc 2.7 program [28], to prepare a file for the analysis in MEGA7.0.14 [29]. Phylogenetic trees were obtained with different methods (MP – maximum parsimony, NJ – neighbour joining and ML – maximum likelihood) and pairwise nucleotide differences were calculated. Obtained sequences were analysed in with three methods: maximum parsimony (MP), neighbor joining (NJ) and maximum likelihood (ML). Obtained sequences were deposited in GenBank MH718837 for the 18S rDNA sequence and MH722215 for the 28S rDNA.

### Collection and external examination of other ladybird species

Since *Hesperomyces virescens* fungi can develop not only on *H. axyridis*, but also on over 30 other ladybirds [1], we decided to examine other ladybirds collected by the same methods and in the same localities as *H. axyridis*. Four hundred specimens of ladybirds of 31 other species (both native and introduced) have been collected and screened for ectoparasitic fungi. The list of these ladybird species: *Adalia bipunctata* (Linnaeus), *Anisosticta novemdecimpunctata* (Linnaeus), *Calvia decemguttata* (Linnaeus), *Chilocorus bipustulatus* (Linnaeus), *Chilocorus renipustulatus* (Scriba), *Coccinella quinquepunctata* Linnaeus, *Coccinella septempunctata* Linnaeus, *Coccinula quatuordecimpustulata* (Linnaeus), *Cryptolaemus montrouzieri* Mulsant, *Exochomus quadripustulatus* (Linnaeus), *Halyzia sedecimguttata* (Linnaeus), *Harmonia quadripunctata* (Pontoppidan), *Hippodamia variegata* (Goeze), *Lindorus lophanthae* (Blaisdell), *Nephus bipunctatus* (Kugelann), *Parexochomus nigromaculatus* (Goeze), *Propylea quatuordecimpunctata* (Linnaeus), *Psyllobora vigintiduopunctata* (Linnaeus), *Rodolia cardinalis* (Mulsant), *Scymnus frontalis* (Fabricius), *Scymnus haemorrhoidalis* Herbst, *Scymnus interruptus* (Goeze), *Scymnus subvillosus* (Goeze), *Scymnus suturalis* Thunberg, *Serangium montazerii* Fürsch, *Stethorus pusillus* (Herbst), *Subcoccinella vigintiquatuorpunctata* (Linnaeus), *Tytthaspis sedecimpunctata* (Linnaeus), *Vibidia duodecimguttata* (Poda).

## Results

We have found three parasitic species on examined *H. axyridis* adults: *Hesperomyces virescens* (Ascomycota: Laboulbeniales, Laboulbeniaceae), *Parasitylenchus bifurcatus* (Nematoda: Tylenchida Allantonematidae) and unknown species belonging to Mermithida (Nematoda). All examined *H. axyridis* specimens were free of parasitic mite *Coccipolipus hippodamiae*.

### Hesperomyces virescens Thaxt

Characteristic yellowish-greenish thalli of *Hesperomyces virescens* were detected on the host integument (Fig 2). Their morphology corresponds to the detailed description by De Kesel [30]. Identification was confirmed by mycologist E.Yu. Blagoveshchenskaya.

**Fig 2.**
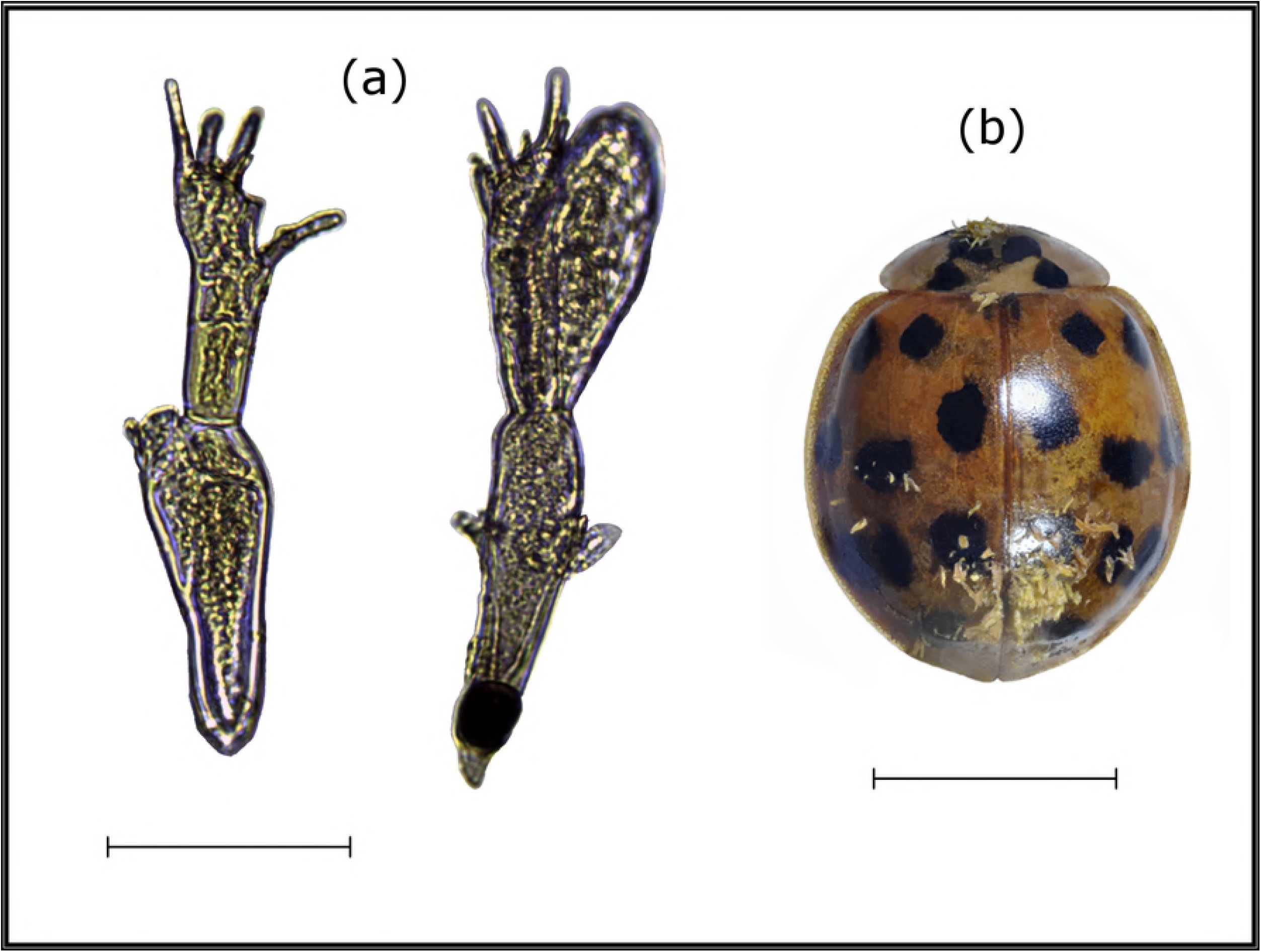
*Hesperomyces virescens* on *Harmonia axyridis*. (a) thalli (bar 50 μm), (b) ladybird covered with thalli (bar 3 mm).

Specimens of *H. axyridis* infested with *H. virescens* were found in all six localities, where the beetles were examined. The distance between the most western locality (Golovinka) and the most eastern (Veseloe) is more than 60 km. The thalli were found in 112 (53%) adults of *H. axyridis* (Table 1) and were situated on elytra, pronotum, sternites, legs and mouthparts of the beetles. It seems that *H. virescens* does not kill or significantly damage *H. axyridis*, since even the beetles covered with large number of fungi thalli moved actively. Males were infested more often than females (62% and 48% respectively). No signs of ectoparasitic fungi have been detected on 400 examined adults of other ladybird species.

**Table 1.** Infestation of *H. axyridis* with *Hesperomyces virescens* and *Parasitylenchus bifurcatus*.

|  | Number<br>of<br>specimens<br>without<br>parasites | Number of<br>specimens<br>infested with<br>only<br>*Hesperomyces*<br>*virescens* | Number of<br>specimens<br>infested with<br>only<br>*Parasitylenchus*<br>*bifurcatus* | Number of<br>specimens<br>infested with<br>*P. bifurcatus* &<br>*H. virescens* | Total |
| --- | --- | --- | --- | --- | --- |
| Males | 22 | 37 | 1 | 1 | 61 |
| Females | 69 | 67 | 9 | 7 | 152 |
| Total | 91 | 104 | 10 | 8 | 213 |

### *Parasitylenchus bifurcatus* Poinar, Steenberg

Parasitic nematodes of the family Allantonematidae were found in *H. axyridis* 18 specimens collected at three out of six sampling locations (see Fig 1). Three different nematode life cycle stages, including a subsequent generation parasitic female, vermiform (infective) female and male, were found in the sampled beetles. The number of females of the subsequent generation varied from 5 to 32 per a beetle. The number of vermiform nematode specimens varied considerably, sometimes being as high as about a couple of hundred per a beetle.

Morphometric analysis of nematodes by standard body features allowed to identify the nematodes extracted from ladybirds as *Parasitylenchus bifurcatus* Poinar, Steenberg, 2012 (Table 2). The characteristic features of *P. bifurcatus* nematodes were as follows: a straight stylet lacking basal thickenings, a forked tail tip in the vermiform females and juvenile males, spicules straight, wedgeshaped or triangular, with narrow bursa and gubernaculum (Fig 3). We found that subsequent generation of parasitic females also has forked tail tip.

**Table 2.** Morphometric characteristics of the nematode *P. bifurcatus* specimens, isolated from the ladybird *Harmonia axyridis* collected in Sochi.

| Character | Parasitic females of the subsequent generation (n= 5) | Vermiform (infective) females (n= 5) | Males (n= 5) |
| --- | --- | --- | --- |
| Body length, $\mu\text{m}$ | 1118.0 (930.0–1660.0) | 590.0 (530.0–670.0) | 403.0 (396.0–480.0) |
| Body width, $\mu\text{m}$ | 124.0 (85.0–184.0) | 12.6 (12.0–13.0) | 15.0(14.0–16.0) |
| Stylet length, $\mu\text{m}$ | | 11.5 (11.0–12.0) | 9.0 (8.0–11.0) |
| Head to excretory pore distance, $\mu\text{m}$ | 155.4 (141.0–165.0) | 50.6 (42.0–57.0) | 66.0(62.0–75.0) |
| Vulva position, % | 89.4 (88.0–93.0) | 88.0 (87.0–90.0) |  |
| Tail length, $\mu\text{m}$ | 39 (25.0–48.0) | 33.2 (30.0–37.0) | 34.4(25.5–40.0) |
| Spicule length, $\mu\text{m}$ | | | 12(11–13) |

**Fig 3.**
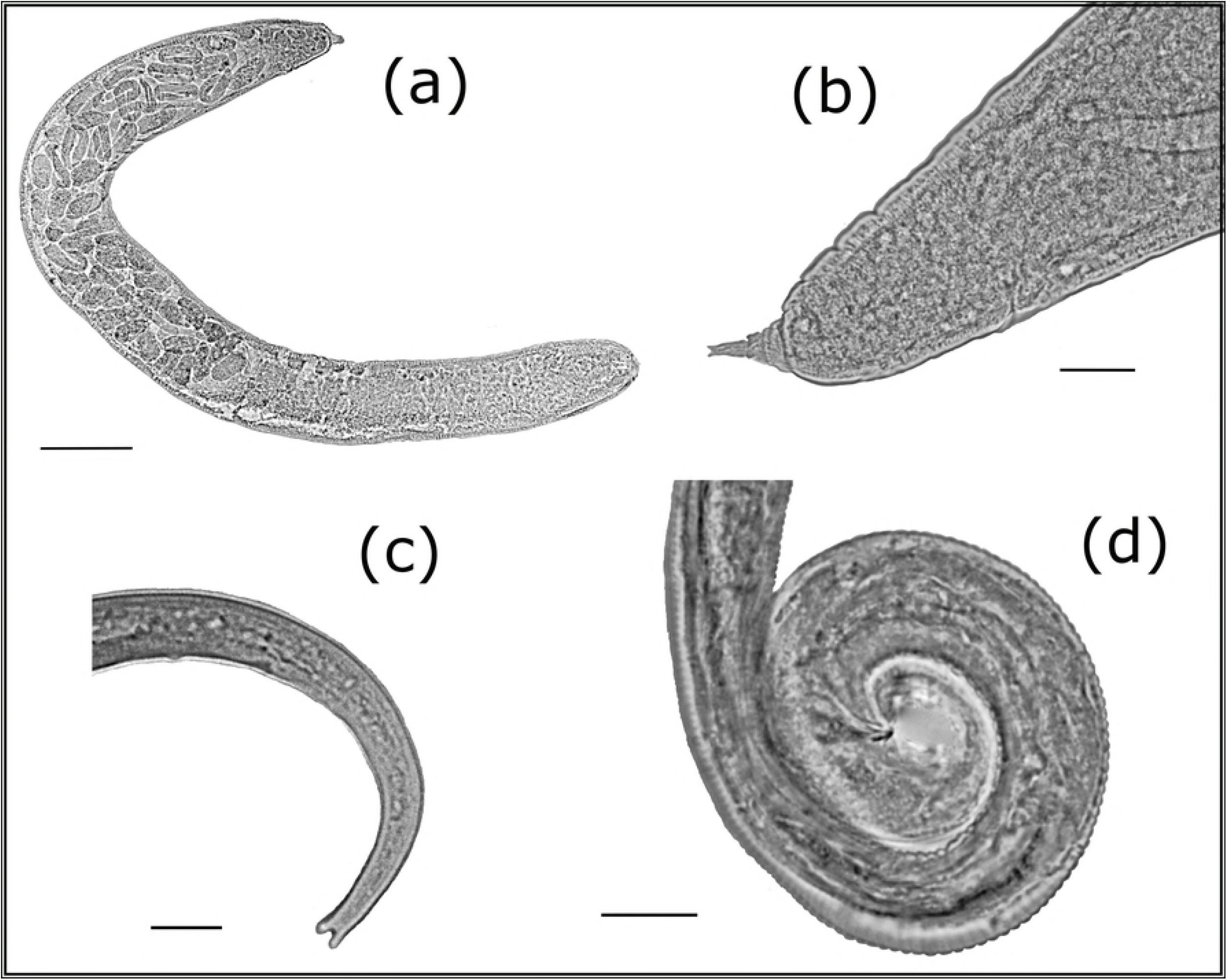
*Parasitylenchus bifurcatus:* (a) subsequent generation female of *P.bifurcatus* (bar 98 μm), (b) subsequent generation female, tail (bar 28 μm), (c) vermiform (infective) female, (d) tail (bar 12 μm), (d) male, tail (bar 12 μm).

Parasitic females of the subsequent generation, sampled in Sochi, were close in the size of their body to the specimens of *P. bifurcates* from Denmark [19]. Their body length of the former was 1118.0 μm (930.0–1660.0) and the body width was 124 μm (85.0–184.0), compared with 1300 μm (920–1600) and 195 μm (158–271) respectively in Danish specimens. In the population of *P. bifurcates*, described in Slovenia [21], females of the subsequent generation were smaller: 886.4 μm (782.0–1098.0) in length and 72.3 μm (59.0–81.0) in width, compared with females from the Sochi population.

The BLAST-search of similar nucleotide sequences in NCBI GenBank was performed for all three obtained sequences. These were the sequences of different clones and isolates of *Parasitylenchus bifurcatus*, which were found as the closest to the obtained 18S rDNA sequence of nematodes from *Harmonia axyridis* ladybirds from Russian Caucasus. All the similar sequences detected by BLAST-search were downloaded and used for comparison. Under all methods of analysis obtained sequence was a member of strongly supported clade (100% bootstrap support) consisting of *P. bifurcatus* sequences plus a sequence of unidentified Allantonematidae (Fig 4). Obtained sequence of *H. axyridis* parasite from Russian Caucaus was 100% identical to some published sequences of *P. bifurcatus* (e.g. clones ‘3l4j’, ‘3j4i’ and ‘PaTyBif1’). Remarkably, an unidentified allantonematid nematode, the sequence of which is deposited as JQ941710 was found to be identical to that obtained in the course of our study was found in 2010 in Germany in the haemocoel of *H. axyridis* (E.L. Rhule, unpublished). It seems, that these nematodes from Germany also belong to the species *P*. *bifurcatus*. All other known 18 rDNA sequences of *P. bifurcatus* differ from our sequence in one or two nucleotides. Several clades demonstrated close, but not securely resolved relationships with the clade containing sequences of *P. bifurcatus*. Under all methods of analysis the 18S rDNA sequence of *Howardula phyllotretae* Oldham, 1933 is in sister relationships with *P. bifurcatus* clade (Fig 4). One such clade consists of the sequences of parasitic tylenchids of fleas: *Rubzovinema* Slobodyanyuk, 1991, *Spilotylenchus* Launay, Deunff & Bain, 1983 and *Psyllotylenchus* Poinar & Nelson, 1973 (Fig 4). Another related clade was represented by species of *Deladenus* Thorne, 1941 and unidentified Tylenchomorpha gen.sp. Other sequences of *Howardula* Cobb, 1921 were most distant from *P. bifurcatus* (different in 59–60 bp) in our analysis and served as a root for obtained cladograms.

**Fig 4.**
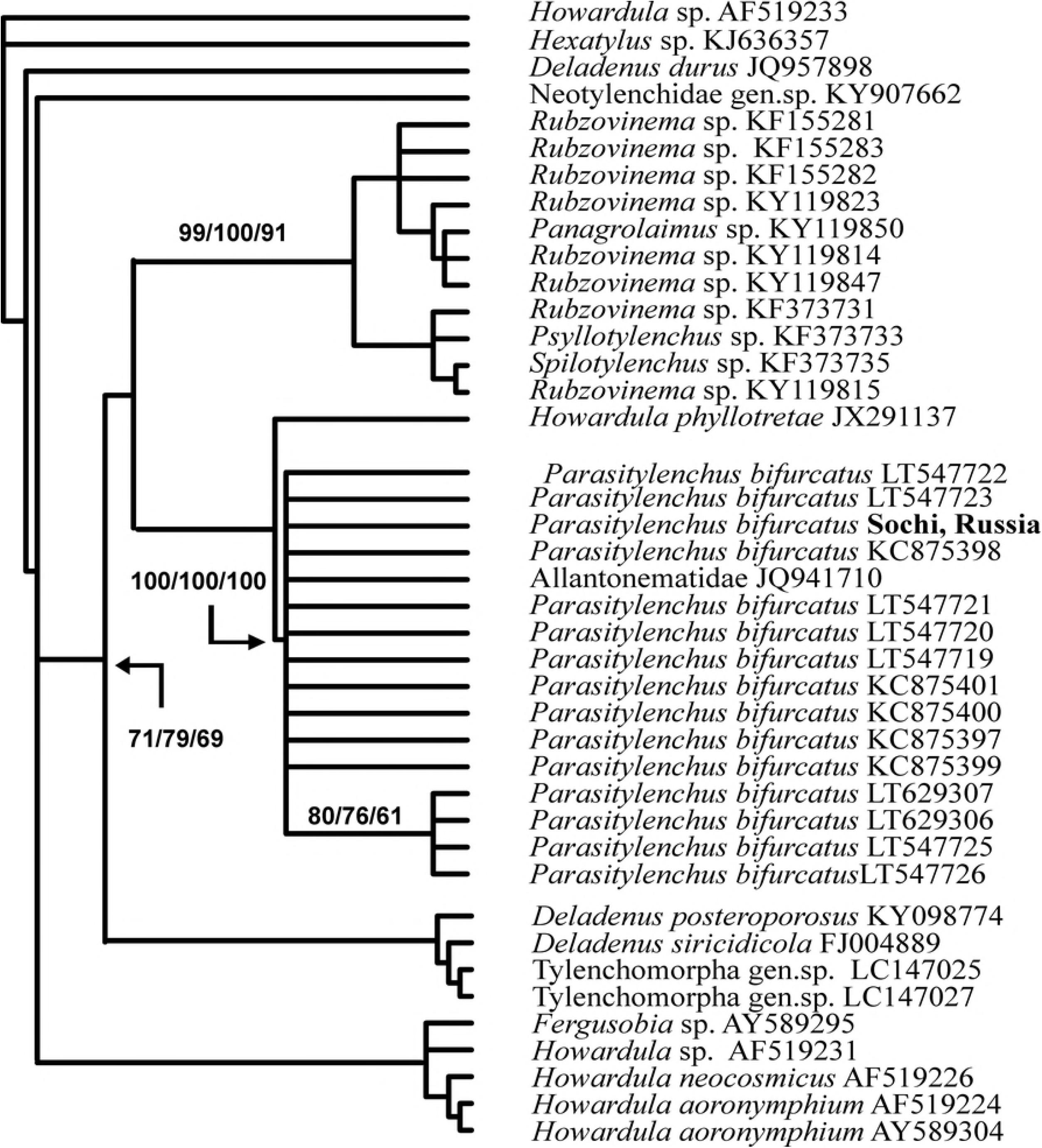
The relationships of *Parasitylenchus bifurcatus* from Sochi, Russia with other groups of insect-associated tylenchids inferred from analysis of partial 18S rDNA. Bootstrap support is given near corresponding nodes in the format MP/NJ/ML.

The known and deposited in NCBI GenBank 28S rDNA sequences of entomoparasitic tylenchids are less numerous than 18S rDNA data. Obtained cladogram demonstrates close relationships of studied nematode with two other sequences obtained for unidentified species of *Parasitylenchus:* DQ328729 and KM245038 (Fig 5). Both are related to the parasites of bark beetles in Russia and Czech Republic, correspondingly. In the level of nucleotide differences these two *Parasitylenchus* sequences are the closest to nematodes from *H. axyridis* (87 and 90 bp), when the nucleotide differences with all other studied entomoparasitic tylenchids exceed 100 bp. The sequence of *Howardula phyllotretae* together with that of *Anguillonema amolensis* Mobasseri, Pedram et Pourjam, 2017 are forming a clade which is in sister position to *Parasitylenchus* Micoletzky, 1922 clade under all methods of analysis (Fig 5). As in 18S rDNA cladogram the sequences of flea parasites (*Rubzovinema, Spylotylenchus, Psyllotylenchus)* are forming well supported clade (Fig 5). The relationships of this latter clade with *Parasitylenchus (Howardula phyllotretae* + *Anguillonema amolense)* clade are strongly supported.

**Fig 5.**
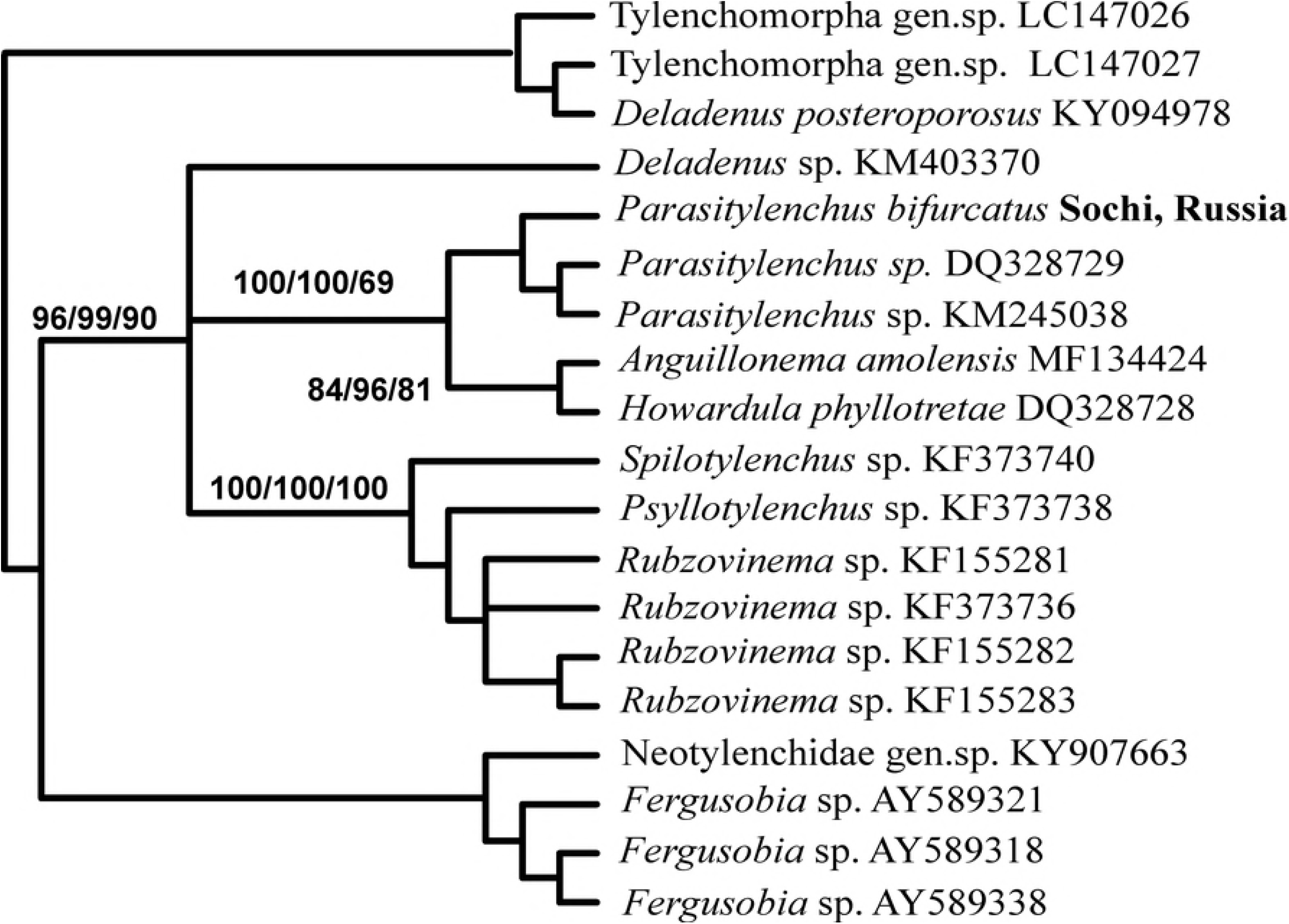
The relationships of *Parasitylenchus bifurcatus* from Sochi, Russia with other groups of insect-associated tylenchids inferred from analysis of partial 28S rDNA. Bootstrap support is given near corresponding nodes in the format MP/NJ/ML.

*Parasitylenchus bifurcatus* was found in 8% of all dissected adults of *H. axyridis*. Prevalence in females and males was 10% and 3% respectively, but this difference is not reliable because of small number of infested specimens. The incidence of the ladybird infestation by *Parasitylenchus bifurcatus* in Sochi is lower, compared with the results of the former studies in the countries of Europe: the incidence of infested ladybirds was as high as up to 35% in Denmark [19], and even up to 47% in Czech Republic [1]. *Parasitylenchus bifurcatus* was found also in specimens infested with *Hesperomyces virescens* and free of fungi, and no correlation between infestation of specimens with *Parasitylenchus bifurcatus* and *Hesperomyces virescens* was observed (Fig 6).

**Fig 6.**
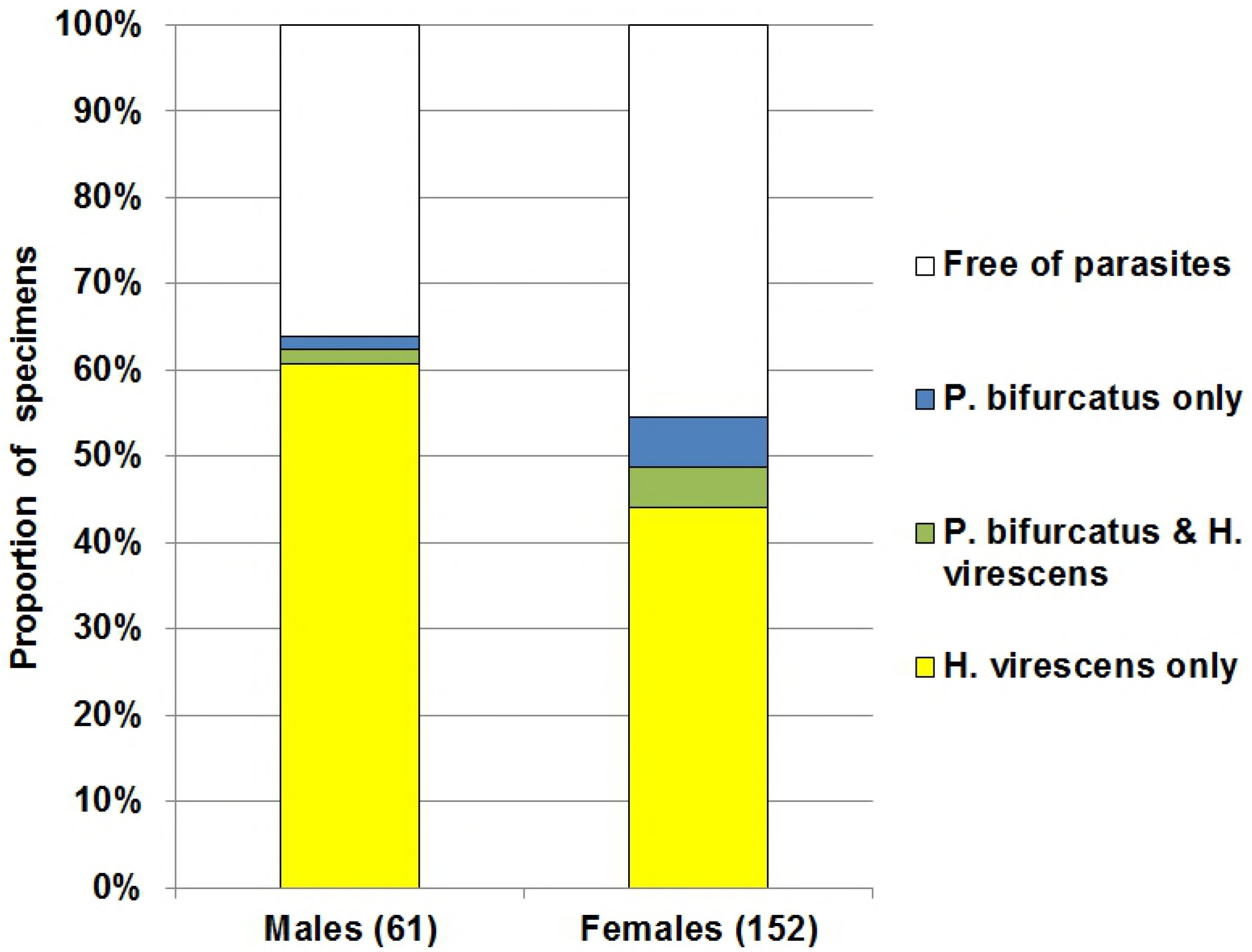
Proportion of *Harmonia axyridis* adults infested with *Hesperomyces virescens* and *Parasitylenchus bifurcatus*. Number of examined specimens is indicated in brackets.

The only one specimen of a nematode, belonging to the order Mermithida, was found in the only ladybird specimen sampled in Veseloe. This was the first documented case of the ladybird H. axyridis infestation by a parasitic nematode of this order in nature. Formerly von Linstow [31] reported *Mermis nigrescens* Dujardin (Mermithida: Mermithidae) to be a parasite of the other ladybird species, *C. septempunctata*.

The infestation of *Adonia variegata, C. septempunctata* und *Semiadalia undecimnotata* with unidentified mermithids in South-East of France was reported by G. Iperti [32]. H. Kaiser and W.R. Nickle [33] described the *Coccinella septempunctata* infestation with *Hexamermis* sp. in Styria, Austria. An overview of reports of coccinellid infestation with mermithid nematodes was presented by G.O. Poinar Jr [34]. A study, specifically focused on the search of parasitic nematodes of this order in the Sochi area, their identification to the species, and determination of the incidence of ladybird infestation is planned for future.

## Discussion

Haelewaters et al. [1] supposed that infection of young populations of *H. axyridis* in some newly invaded regions by *H. virescens* and *P. bifurcatus* (North America, Netherlands), occurred as a result of acquisition of native natural enemies. In the Caucasus co-invasion hypothesis is more relevant than acquisition hypothesis. First, *H. virescens* and *P. bifurcatus* were absent in the region and appeared at the same time with their host. Locality of the current records of the parasites (Sochi) is situated at more than 1400 km from the nearest known localities of both parasitic species. Second, other examined ladybird species are not infested with *H. virescens* in the Caucasus. Other ladybirds were not dissected to find nematodes, but, since *H. axyridis* is the only known host of *P. bifurcatus*, there is no reason to suggest an *H. axyridis* got infected from other ladybird species. Third, an analysis of the phylogenetic relationships of *Parasitylenchus bifurcatus* based on 18S rDNA demonstrated complete identity of 18S rDNA sequence of these nematodes from Russian Caucasus with some strains or clones found in Western and Southern Europe. Cautiously, such identity can be considered as an indication on the possible transfer of parasites together with insect hosts from Western part of Eurasia.

Since *H. axyridis* was released for biological control of pests in the Caucasus, it was unclear, how the population of *H. axyridis* appeared: as a result of these releases or as a result of expansion of its European range [11, 16]. The study of parasites has shed light on this question. Both *H. virescens* and *P. bifurcatus* affect only adults, do not occur on other life stages and exclusively spread by activities of the host [18]. Transmission takes place only from adult to adult, therefore the direct contact between different generations of beetles is necessary for to maintain the life cycle of the parasites. Since different generations are kept separately in laboratory culture [35], it is free of parasites. Therefore, detection of *H. virescens* and *P. bifurcatus* indicates that population of *H. axyridis* in the Caucasus cannot derive exclusively from specimens released from laboratory culture. At least part of ancestors of the Caucasian population of *Harmonia axyridis* are from European invasive range. On the other hand, admixture of released specimens is not excluded, since *H. axyridis* was released in Sochi for several decades. The complex invasion scenaria are common place for alien insects in general [36] and for *Harmonia axyridis* in particular [37]. This case and some other recent studies [2] confirm that parasitological analysis is a promising approach of revealing of invasion rotes.

Roy et al [17] supposed that connection of *H. virescens* with *H. axyridis* coupled with the rapid expansion of *H. axyridis* globally suggests that this parasite will continue to spread throughout the rest of the world. Spread of *H. virescens* to the Caucasus confirms this suggestion.

*Hesperomyces virescens* is found to be widespread and common on *H. axyridis* at the Black sea coast of the Caucasus. But in spite of high prevalence of *H. virescens* on *Harmonia axyridis* we have not found it on other potential hosts in the Caucasus. No signs of Laboulbeniales ectoparasites have been detected on other ladybird species, in spite seven of them were indicated by Ceryngier and Twardowska [38] as hosts of *Hesperomyces virescens* in other regions: *Adalia bipunctata, Chilocorus bipustulatus, Chilocorus renipustulatus, Coccinula quatuordecimpustulata, Propylea quatuordecimpunctata, Psyllobora vigintiduopunctata* and *Tytthaspis sedecimpunctata*. The same situation was previously observed by A. De Kesel in Europe in spite some potential hosts overwintered at the same sited with *Harmonia axyridis* [1]. Also Cottrell and Riddick [18] found reduced interspecific transmission of *H. virescens* (under laboratory conditions) and hence suggested the existence of hostadapted isolates or strains of *H. virescens*.

*Hesperomyces virescens* does not seriously damage its hosts [17], but *Parasitylenchus bifurcatus* capable of causing significant harm [19]. Co-infection of *H. axyridis* with *H. virescens* and *P. bifurcatus* was recorded in Netherlands and positive association between these parasites was detected that correlated with a reduced number of live beetles [20]. No correlation between infestation by *H. virescens* and *P. bifurcatus* has been yet recorded in the Caucasus. But the number of collected specimens was small, so possibility of such correlation cannot be ruled out. The study of co-infection of *H. axyridis* by these two parasites is an intriguing subject for future studies, since co-infections might result in lower survival rates.

The obtained nucleotide sequences are significant as the confirmation of primary parasitic nematode identification based on morphological features. Several sequences for 18S rDNA of *P. bifurcatus* are deposited in NCBI GenBank, and newly obtained ones for the specimens from Sochi are identicval with some of these. Unlike 18S rDNA data, those for large ribosomal subunit (28S rDNA) of *Parasitylenchus* nematodes are quite scarce and can only prove the sufficient informative value of this locus for phylogenetic studies of entomoparastic tylenchids. These data securely demonstrated the clustering of *P. bifurcatus* sequence with two known sequences of *Parasitylenchus* nematodes. The results of both 18S and 28S analyses revealed some incongruence in contemporary taxonomy of these nematodes. Thus, it is obvious in the obtained phylogenetic trees that some genera are polyphyletic: we can find the species of *Howardula* in three clades of 18S rDNA cladogram, *Deladenus* sequences in two clades of 18S and 28S rDNA cladograms.

## Conclusions

1. The population of *H. axyridis* in the Caucasus recently invaded by this species is infested with two parasite species, which are recorded for the Caucasus and Russia for the first time: *Hesperomyces vinirescens* and *Parasitylenchus bifurcatus*. Probably these parasites have appeared in the region as a result of co-invasion with *H. axyridis*.
2. Population of *H. axyridis* in the Caucasus appeared as a result of expansions of European range, as 18S rDNA sequences of Caucasian *Parasitylenchus bifurcatus* and those from Western Europe are 100% identical. It cannot derive exclusively from specimens released for biological control of pests, because laboratory cultures are free of these parasites.
3. Though *Hesperomyces vinirescens* develops on many ladybird species in other regions, its only known host in the Caucasus is *Harmonia axyridis*.
4. An unidentified species of the order Mermithida is recorded on *Harmonia axyridis* in the Caucasus. It is the first documented case of the ladybird *H. axyridis* infestation by a parasitic nematode of this order in nature.

## Acknowledgements

We are grateful to E.Yu. Blagoveshchenskaya (Department of Mycology and Algology of Moscow State University) for identification of the parasitic fungi and taking the photo of thalli, to P. Ceryngier (Faculty of Biology and Environmental Sciences, Cardinal Stefan Wyszyński University) for valuable information and to T.A. Mogilevich for the help in screening of the ladybirds for parasites.

## Supporting information captions

S1 Table. Localities of *Harmonia axyridis* and its parasites *Hesperomyces virescens* and *Parasitylenchus bifurcatus* in the Caucasus

## References

1. Haelewaters D, Zhao SY, Clusella-Trullas S, Cottrell TE, De Kesel A, Fiedler L, et al. Parasites of *Harmonia axyridis*: current research and perspectives. Biocontrol (Dordr). 2017;62: 355–371. doi: 10.1007/s10526-016-9766-8.

2. Reshetnikov AN, Sokolov SG, Protasova EN. Detection of a neglected introduction event of the invasive fish *Perccottus glenii* using parasitological analysis. Hydrobiologia. 2017;788(1): 65–73. doi: 10.1007/s10750-016-2987-0.

3. Vilcinskas A, Stoecker K, Schmidtberg H, Röhrich CR, Vogel H. Invasive harlequin ladybird carries biological weapons against native competitors. Science. 2013;340(6134): 862–863. doi: 10.1126/science.1234032.

4. Orlova-Bienkowskaja MJ, Ukrainsky AS, Brown PMJ. *Harmonia axyridis* (Coleoptera: Coccinellidae) in Asia: a re-examination of the native range and invasion to southeastern Kazakhstan and Kyrgyzstan. Biol Invasions. 2015;17(7): 1941–1948. doi: 10.1007/s10530-015-0848-9.

5. Roy HE, Brown PMJ, Adriaens T, Berkvens N, Borges I, Clusella-Trullas S, et al. The harlequin ladybird, *Harmonia axyridis*: global perspectives on invasion history and ecology. Biol Invasions. 2016;18(4): 997–1044. doi: 10.1007/s10530-016-1077-6.

6. Roy HE, Adriaens T, Isaac NJ, Kenis M, Onkelinx T, Martin GS, et al. Invasive alien predator causes rapid declines of native European ladybirds. Divers Distrib. 2012;18(7): 717–725. doi: 10.1111/j.1472-4642.2012.00883.x.

7. Ukrainsky AS, Orlova-Bienkowskaja MJ Expansion of *Harmonia axyridis* Pallas (Coleoptera: Coccinellidae) to European Russia and adjacent regions. Biol Invasions. 2014;16(5): 1003–1008 doi: 10.1007/s10530-013-0571-3.

8. Goryacheva I, Blekhman A, Andrianov B, Romanov D, Zakharov I. *Spiroplasma* infection in *Harmonia axyridis*-Diversity and multiple infection. PLoS One. 2018;13(5): e0198190. doi: 10.1371/journal.pone.0198190.

9. Kuznetsov VN. Far Eastern coccinellids in the Transcaucasia. Zashchita rasteniy. 1988;5: 19. [In Russian]

10. Mogilevich TA. My experiments with the ladybird *Harmonia axyridis* [cited 6 August 2018]. In: Beetles and coleopterologists 2012. Available from: https://www.zin.ru/aNIMAliA/Coleoptera/rus/mogilev1.htm. [in Russian]

11. Belyakova NA, Reznik SY. First record of the harlequin ladybird, *Harmonia axyridis* (Coleoptera: Coccinellidae) in the Caucasus. Eur J Entomol. 2013;110(4): 699–702.

12. Orlova-Bienkowskaja MJ, Mogilevich TA. The first record of Asian ladybird *Harmonia axyridis* (Pallas, 1773) (Coleoptera: Coccinellidae) in Kabardino-Balkaria and the history of the expansion of this alien species in the Caucasus and south of European Russia in 2002–2015. Caucasian Entomological Bulletin. 2016;12(1): 93–98. [In Russian]

13. Alekseev SK, Butaeva FG. Ladybirds (Coleoptera: Coccinellidae) of the vicinity of Alagir City (Republic of North Ossetia-Alania). In: Actual problems of chemistry, biology and biotechnology: Proceedings of the 10th All-Russian Scientific Conference (11–13 May 2016). North Ossetia State Univeristy. Vladikavkaz: Publishing house of North Ossetia State Uiversity; 2016. pp. 71–75. [in Russian]

14. Kalashian MY, Ghrejyan TL, Karagyan GH. Harlequin ladybird *Harmonia axyridis* Pall. (Coleoptera, Coccinellidae) in Armenia. Russ J Biol Invasions. 2017;8(4): 313–315. doi: 10.1134/S207511171704004X.

15. Bugaeva LN, Ignat’eva TN, Novikov YuP., Kashutina EV. Problem of protection of vegetables at organic food agriculture. Newsletter of the East Palaearctic Regional Section of the Internationalorganizations for the biological control of harmful animals and plants, 2011;42: 32–35. [in Russian]

16. Orlova-Bienkowskaja MJ. The outbreak of harlequin ladybird *Harmonia axyridis* (Pallas, 1773) (Coleoptera, Coccinellidae) in the Caucasus and Possible Sources of Invasion. Russ J Biol Invasions. 2014;5(4): 275–281. doi: 10.1134/S2075111714040055.

17. Roy HE, Rhule E, Harding S, Handley LJL, Poland RL, Riddick EW, Steenberg T. Living with the enemy: parasites and pathogens of the ladybird *Harmonia xyridis*. Biocontrol (Dordr). 2011;56(4): 663–679. doi: 10.1007/s10526-011-9387-1.

18. Cottrell TE, Riddick EW. Limited transmission of the ectoparasitic fungus *Hesperomyces virescens* between lady beetles. Psyche. 2012;814378. doi: 10.1155/2012/814378.

19. Poinar GO, Steenberg T. *Parasitylenchus bifurcatus* n. sp.(Tylenchida: Allantonematidae) parasitizing *Harmonia axyridis* (Coleoptera: Coccinellidae). Parasit Vectors. 2012;5(1): 218. doi: 10.1186/1756-3305-5-218.

20. Raak-van den Berg CL, van Wielink PS, de Jong PW, Gort G, Haelewaters D, Helder J, Karssen G, van Lenteren JC. Invasive alien species under attack: natural enemies of *Harmonia axyridis* in the Netherlands. Biocontrol (Dordr). 2014;59: 229–240. doi: 10.1007/s10526-014-9561-3.

21. Gerič Stare B, Širca S, Urek G. First report of nematodes *Parasitylenchus bifurcatus* Poinar & Steenberg, 2012 parasitizing multicolored Asian lady beetle *Harmonia axyridis* (Pallas, 1773) in Slovenia. Acta Agric Slov. 2017;109(2): 457–463. doi: 10.14720/aas.2017.109.2.28.

22. Seinhorst JW. A rapid method for the transfer of nematodes from fixative to anhydrous glycerin. Nematologica. 1959;4(1): 67–69.

23. Holterman M, van der Wurff A, van den Elsen S, van Megen H, Bongers T, Holovachov O, et al. Phylum-wide analysis of SSU rDNA reveals deep phylogenetic relationships among nematodes and accelerated evolution towards crown clades. Mol Biol Evol. 2006;23: 1792–1800. doi: 10.1093/molbev/msl044.

24. Floyd RM, Rogers AD, Lambshead PJD, Smith CR. Nematode-specific PCR primers for the 18S small subunit rRNA gene. Mol Ecol Notes. 2005;5: 611–612. doi: 10.1111/j.1471-8286.2005.01009.x.

25. Nadler SA, Bolotin E, Stock SP Phylogenetic relationships of *Steinernema* Travassos, 1927 (Nematoda: Cephalobina: Steinernematidae) based on nuclear, mitochondrial and morphological data. Systematic Parasitology. 2006;63: 161–181.

26. Thompson JD, Gibson TJ, Plewniak F, Jeanmougin F, Higgins, DG. The Clustal_X windows interface: flexible strategies for multiple sequence alignment aided by quality analysis tools. Nucleic Acids Res. 1997;24: 4876–4882. doi: 10.1093/nar/25.24.4876.

27. Altschul SF, Gish W, Miller W, Myers EW, Lipman DJ. Basic local alignment search tool. J Mol Biol. 1990;215: 403–410.

28. Nicholas KB, Nicholas HB Jr., Deerfield DW. Multiple Sequence Alignment Editor and Shading Utility, Version 2.7.000: 1997. [cited 6 August 2018] Available from: http://www.psc.edu/biomed/genedoc.

29. Kumar S, Stecher G, Tamura K. MEGA7: Molecular evolutionary genetics analysis. Version 7.0 for bigger datasets. Mol Biol Evol. 2016;33: 1870–1874. doi: 10.1093/molbev/msw054.

30. De Kesel A. *Hesperomyces* (Laboulbeniales) and coccinellid hosts. Sterbeeckia. 2011;30: 32–37.

31. Linstow OFP von. Das Genus *Mermis*. Archiv für mikroskopische Anatomie und Entwicklungsmechanik 1898;53: 149–168.

32. Iperti G. Les parasites des Coccinelles aphidiphages dans les Alpes-Maritimes et les Basses-Alpes. Entomophaga. 1964;9: 153–180.

33. Kaiser H, Nickle WR. Mermithiden (Mermithidae, Nematoda) parasitieren Marienkafer (*Coccinella septempunctata* L.) in der Steiermark. Mitteilungen Naturwissenschaftlicher Verein für Steiermark. 1985: 115–118.

34. Poinar GO Nematodes for biological control of insects. Boca Raton: CRC Press; 1979.

35. Izhevsky SS. Introduction and application of entomophages. Moscow: Agropromizdat; 1990. [In Russian]

36. Garnas JR, Auger-Rozenberg MA, Roques A, Bertelsmeier C, Wingfield MJ, Saccaggi DL, Roy HE, Slippers B. Complex patterns of global spread in invasive insects: eco-evolutionary and management consequences. Biol Invasions. 2016;18(4): 935–952. doi: 10.1007/s10530-016-1082-9.

37. Lombaert E, Guillemaud T, Lundgren J, Koch R, Facon B, Grez, A, et al. Complementarity of statistical treatments to reconstruct worldwide routes of invasion: the case of the Asian ladybird *Harmonia axyridis*. Mol Ecol. 2014;23(24): 5979–5997. doi: 10.1111/mec.12989.

38. Ceryngier P, Twardowska K. *Harmonia axyridis* (Coleoptera: Coccinellidae) as a host of the parasitic fungus *Hesperomyces virescens* (Ascomycota: Laboulbeniales, Laboulbeniaceae): A case report and short review. Eur J Entomol. 2013;110(4): 549–557.

